# Scale Bar of Aging Trajectories for Screening Personal Rejuvenation Treatments

**DOI:** 10.1101/2022.01.17.476558

**Authors:** Xilin Shen, Bingbing Wu, Wei Jiang, Yu Li, Yuping Zhang, Kun Zhao, Nanfang Nie, Lin Gong, Yixiao Liu, Xiaohui Zou, Jian Liu, Jingfen Jin, HongWei Ouyang

## Abstract

Although aging is an increasingly severe healthy, economic, and social global problem, it is far from well-modeling aging due to the aging process’s complexity. To promote the aging modeling, here we did the quantitative measurement based on aging blood transcriptome. Specifically, the aging blood transcriptome landscape was constructed through ensemble modeling in a cohort of 505 people, and 1138 age-related genes were identified. To assess the aging rate in the linear dimension of aging, we constructed a simplified linear aging clock, which distinguished fast-aging and slow-aging populations and showed the differences in the composition of immune cells. Meanwhile, the non-linear dimension of aging revealed the transcriptome fluctuations with a crest around the age of 40 and showed that this crest came earlier and was more vigorous in the fast-aging population. Moreover, the aging clock was applied to evaluate the rejuvenation effect of molecules *in vitro*, such as Nicotinamide Mononucleotide (NMN) and Metformin. In sum, this study developed a *de novo* aging clock to evaluate age-dependent precise medicine by revealing its fluctuation nature based on comprehensively mining the aging blood transcriptome, promoting the development of personal aging monitoring and anti-aging therapies.

## Introduction

Life expectancy has increased dramatically in the past 150 years. It is expected that the 1.5 billion people aged 65 years or over worldwide will outnumber adolescents and youth aged 15 to 24 years (1.3 billion) in 2050^1^. People aged 65 years and older are experiencing the aging process, characterized by progressive impairment and loss of physiological integrity and function, leading to an increased vulnerability to death^2^. Therefore, the world is facing an aging challenge.

Aging, a complex biological process, is far from well modeled though significant efforts have been put into understanding the aging process and revealing patterns in immune-aging^3^ and inflammatory-aging^4^ perspectives. Until now, ‘Omics’ technologies *(e.g*., genomics, metabolomics, metagenomics, proteomics, and transcriptomics) have been widely applied to investigate and model the aging process^5^. Among these Omics, transcriptomics by RNA sequencing is a mature and relatively low-cost omics technology and has already been in clinical use. In addition, transcriptome-based aging clocks, including the analyses of peripheral blood mononuclear cells (PBMCs)^6^, muscle^7^, and dermal fibroblast^8^, are high in interpretability without compromising accuracy^9^ compared with other aging clocks. However, most studies modeled aging as a static linear process^6–8^, failure to model it as a dynamic process^10^. Given that recent studies have shown the diversified early aging signs or pace^11^ at middle age and the fluctuation in plasma protein level^10^, examining the transcriptome changes of blood samples in midlife can help investigate and model the aging process.

In the search for anti-aging intervention and drugs, a quantitative measurement of sample biological age before and after intervention cannot be achieved without accurate modeling. However, the lack of accuracy prevented their scientific and clinical usage of the aging clocks. Of note, the application of transcriptome-based aging clock in drug anti-aging effect assessment is still absent, leaving a gap between model construction and application. Therefore, an accurate and applicable transcriptome-based aging clock is required. This study aims to construct the aging trajectories using blood transcriptomics and successfully developed a new aging clock capable of reflexing the linear and dynamic changes with high accuracy using ensemble modeling. Moreover, we investigated the possibility of using the new aging clock to screen rejuvenation treatments.

## Results

### 1. Trajectories of Aging Gene Expression Form Functional Modules

To dissect the transcriptome landscape of the aging process, we did the HiseqX sequencing on blood samples from a cohort of 505 volunteers, including 208 male and 297 female participants with the age range from 18 to 68, with a median of 36 (Fig. S1-A). First, we grouped genes with similar trajectories by unsupervised hierarchical clustering to identify the changing pattern of age-related genes. Eight modules were identified, of which five (Clusters 1-5) showed an upward trend, and Clusters 6-8 had downward patterns (Fig. 1-A, B). As visualized in trajectory bundles (Fig. 1-B), some patterns were generally linear, but others were non-linear. In some of the modules (Clusters 5-8), gene expressions changed steadily, while other trajectories indicated dramatic changes in a specific age range. Gene Ontology (GO) Enrichment analysis was then conducted to infer its related biological function. The dot plot showed top enriched GO terms in each module (Fig. 1-C, Supplement table 1). The first module expression was enhanced at the age of 25-35, and its genes are related to ubiquitin activity and immune cell proliferation. The second module was wave-like, and the related genes in this module regulate transcription factor complex and interleukin-8 secretion. The age of 45 is the boosting point for the third module expression, of whose genes were associated with mitochondria activity. The expression of the fourth and fifth modules, including the genes enriched in neutrophil immune activity, was increased at the age of 35-45. The other three modules (Clusters 6-8) with downward trends were mainly involved in translation, including that the top terms were protein targeting to membrane, RNA helicase activity, and viral translation, respectively. These biological processes, enriched in these modules, correspond to previous studies of ubiquitin^12^, immune cell^13^, mitochondria^14^, ribosome^15^ in aging. In sum, we mapped the trajectories of the expression pattern of aging-related gene expression.

**Figure 1.**
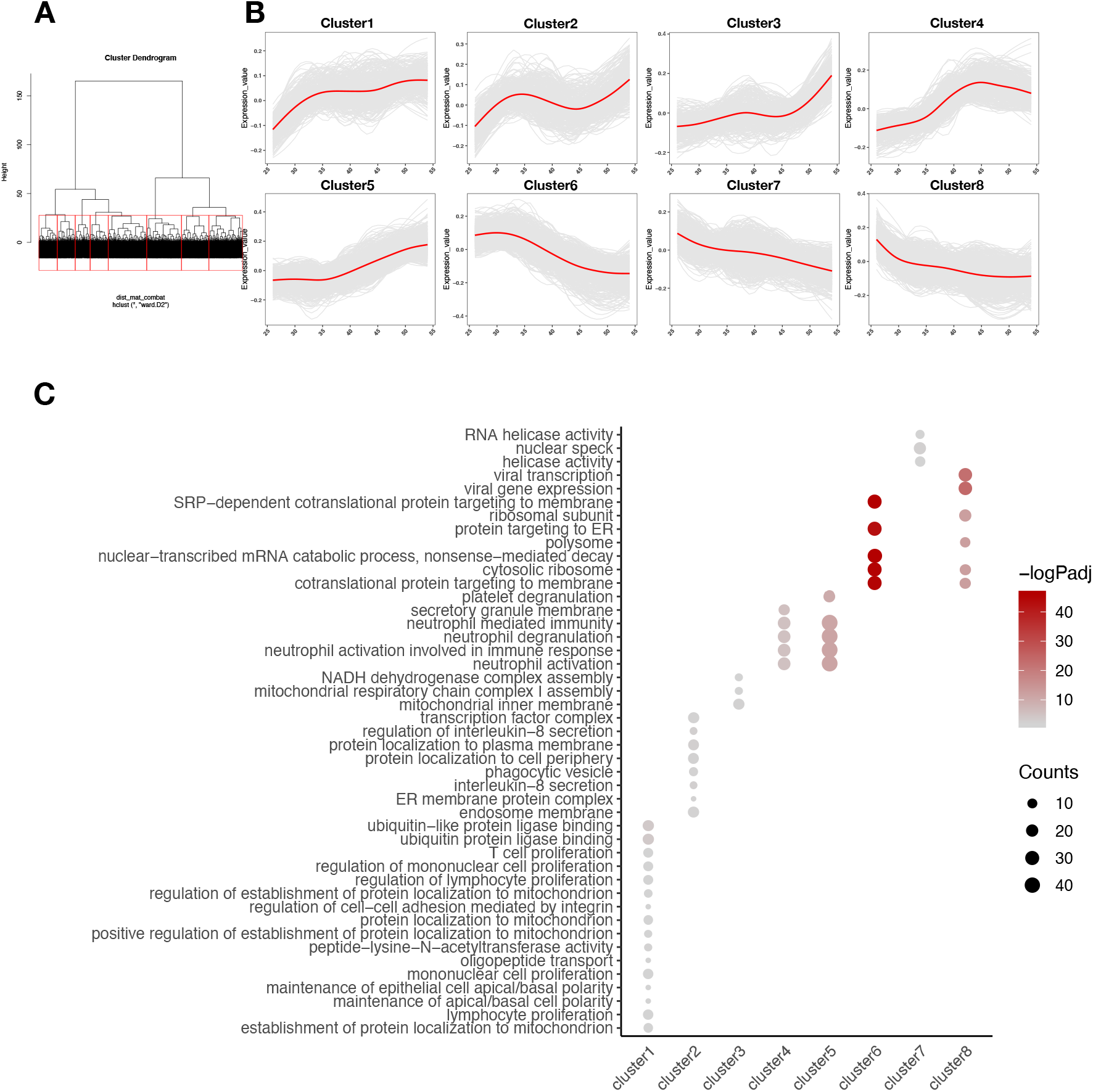
Trajectories of gene expression throughout age form functional modules. (A) Hierarchical clustering of gene expression trajectories. A red box highlighted each cluster. (B) Eight clusters of the aging pattern (five up-regulated, three down-regulated, respectively). Redline was indicating the fitting curve created by loess. (C) Top GO terms in which each aging pattern is involved. Color showing the p adjusted value of enrichment analysis. Dot size showing the number of genes hit by GO terms.

### 2. Identification of Linear Age-Related Genes (ARGs)

Linear fitting was first applied to identify Linear ARGs. As females have a longer lifespan than man^16^, we applied the linear fitting for each gene with age and gender as variables. 1,138 genes significantly affected by age were identified as Linear ARGs (t-test for age effect: p-value < 0.05). Five hundred thirty genes were downregulated, and 608 were upregulated considering the age effect (Supplement table 2). *FMNL1* and *NELL2* belong to the top five Linear ARGs (Fig. 2-A). Consistent with the previous findings, *FMNL1* was reported increased in arterial endothelial aging^17^, and *NELL2* was found to be downregulated in the elderly^13^. 1,221 (81.5%) of all previously summarized ARGs in a meta-analysis^6^ were identified here, including 184 (15.1%) Linear ARGs. 94% of these Linear ARGs were associated with chronological age in the same direction (Figure S2-A).

**Figure 2.**
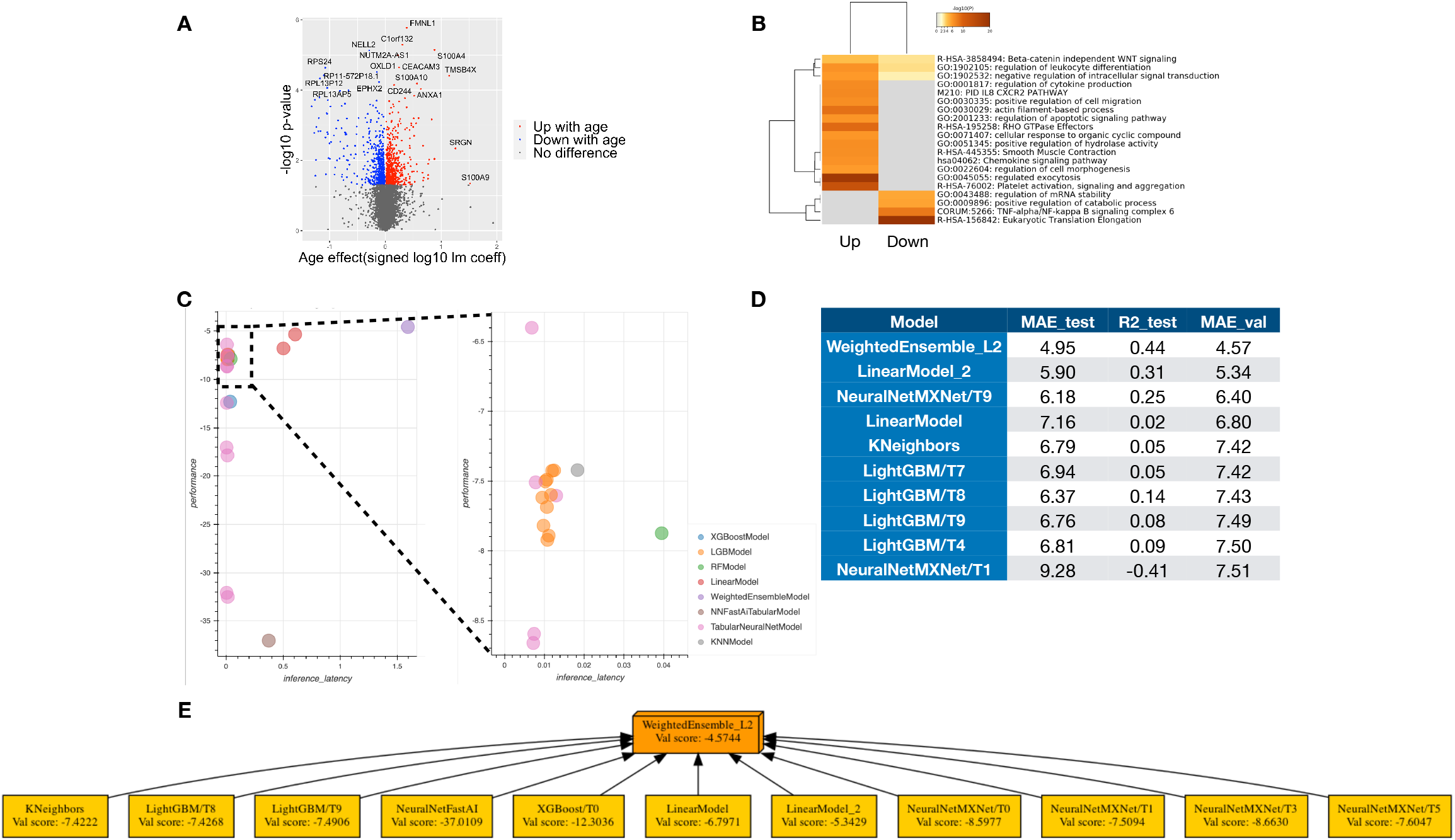
The construction of the ensemble model as the aging clock. (A) Volcano plot shows age-related gene discovered by linear fitting (Linear ARGS). (p-value <0.05, 608 of which up with age and 530 of which down with age) X-axis showed signed log10 of linear fitting coefficient; y-axis showed negative of log10 of the p-value. (B) Heat map of pathway enriched for the age-related gene in GO and Reactome terms. (C) Performance of models constructed by AutoGulon. MAE is for mean absolute error. The X-axis shows time latency for inference. Y-axis shows model performance measured in the MAE validation set. (D) Performance metrics in top10 models. R2: R squared, test: test set, Val: validation set. The whole table is in supplementary data. (E) Models that contribute to ensemble WeightedEnsemble_L2 model. The score is negative MAE, as the algorithm selects the model by the rule that the model with a higher score is better than the control.

Moreover, Metascape^18^ enrichment analyses were performed on Linear ARGs of both directions, respectively. The top enriched terms for upregulated Linear ARGs were platelet activation, signaling and aggregation, regulated exocytosis, and apoptotic signaling pathway. Those downregulated Linear ARGs were Eukaryotic translation elongation, TNF-alpha/NF-kappa B signaling complex, and positive regulation of the catabolic process (Fig. 2-B, Supplement table 3). Of note, the downregulated Linear ARGs, including ribosome genes (*e.g*., 25 RPS-genes and 37 RPL-genes (pseudogene included)), are highly related to the biological processes in translation, similar to the previous results^6^. Among the top Linear ARGs, *RPL5*, *RPL11*, and *RPL23A* were reported as participants of ribosome biogenesis stress followed by the p53 activation^19^.

Furthermore, the percentage of variance explained by sex and age for each gene was computed (Figure S2-B, C). It showed that the age-related genes were also significantly related to sex, such as *RPS4Y1*, encoding a thioredoxin-binding protein, apart from genes encoded by the Y chromosome. Taken together, we identified ARGs by applying linear fitting and Metascape enrichment analyses.

### 3. Ensemble Model as Aging Clock Was Constructed by Auto Machine Learning Framework

To predict the biological age, we constructed an aging clock based on Linear ARGs. The auto machine-learning technique was AutoGluon^20^ by applying hyperparameter search, model selection, and ensemble model construction (See Method). The cohort was first divided as train set and test set with a ratio of 3:1. Then, the train set was further divided, 80% of which was used for model construction and 20% for validation. Finally, the top models were trained and tested in mean absolute error (MAE) (Fig. 2-C, D), and the weighted ensemble model showed the best accuracy in the separated test set (Fig. 2-C, D and Supplement table 4). An ensemble is a collection of models whose predictions are combined by the weighted averaging or voting^21^. In our case, it is constructed from 11 selected models (Fig. 2-E). The feature importance of the weighted ensemble model was measured by permutation (Supplement table 5). *NT5E* (also referred to as *CD73)*, which is among the top ranked features, was reported related to NAD metabolism and calcification of joints and arteries^22^, and *CRLG5*-decreased in the age-related nuclear cataracts^23^.

### 4. Linear Aging Clock Predicts Quick- and Slow-aging Population, Respectively

A simpler model can provide a better interpretation and a lower expected risk in the application. The Linear Model_2 showed second-best accuracy (Fig. 2-D) while its structure was much simpler than the ensembled model (Fig. 2-E). Thus, we applied more tactile parameter searches by elastic-net for a linear model as a substitute for the ensemble model (See Method and FigS3-A & Supplement table 6) and yielded the best model with an MAE of 5.02 and 0.54 in the separated test set (Fig. 3-A, B). The model remained accurate upon the down-sampling of genes. Sampling and the broken-stick test were applied to find a reduced model with fewer genes, which estimated a turning point of 219 genes. A reduced model could achieve an MAE of 5.37 with 200 genes (FigS3-C). This model outperformed the previous blood transcriptome-based aging clock constructed in ribo-minus PBMC^24^ (MAE=5.68) and multiple cohort model constructed in whole-blood gene expression array data^6^ (MAE=7.8), as well as other transcriptome-based aging clocks constructed in muscle gene expression^7^ (MAE=6.24).

**Figure 3.**
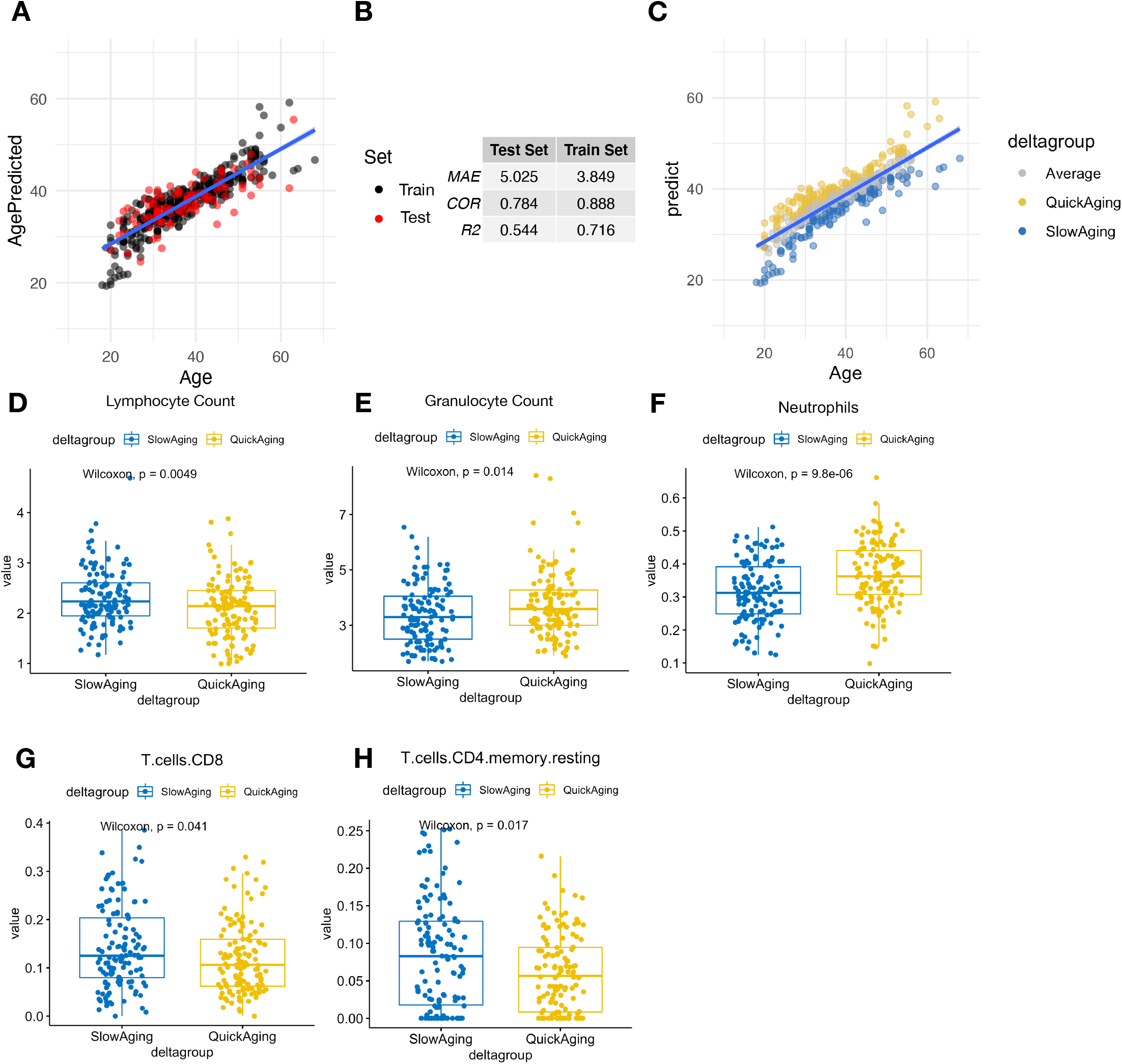
Construction of Biological-meaningful Aging Clock predicted population-based Quick-aging and Slow-aging group. (A) Aging clock constructed visualized with actual age against predicted age. (each dot represents a person, black dots: samples used in training, red dots: samples used in the validation) (B) Performance metrics in the regression model. MAE: mean absolute error, R2: R squared. Correlation: Pearson correlation score. (C) Aging clock colored with “Quick-aging” and “Slow-aging” population defined as delta group. Top indicating predicted Quick-aging with curated delta age in the top quarter of the population, bottom indicating predicted Slow-aging with curated delta age in last quarter of the population. (D)-(I) Box plot shows the contrast of cell fraction calculated by deconvolution(F-I)/ cell count in blood test(D-E) between predicted Quick-aging and Slow-aging population. (Wilcoxon test)

Prediction of the aging clock was used to define the aging rate. As commonly suggested by previous studies, the difference between the model predicted age and chronological age was used to evaluate the personal status of aging^9^. The prediction error distribution was adjusted for age and data set differences (Fig S3-B, D). The population was classified into average, quick-aging, and slow-aging groups (Fig. 3-C). Then we asked if the blood test result and immune cell composition differed in quick-aging and slow-aging populations. As blood test results showed, the slow-aging group showed a significantly higher lymphocyte count (p-value = 0.0049, Wilcoxon test) and significantly lower granulocyte count (p-value = 0.014, Wilcoxon test) (Fig. 3-D, E), indicating a younger blood cell count phenotype^25^. Then we applied Cibersortx^26^, an approach for digital cytometry, and its built-in blood immune cell signature LM22^27^ to deconvolute the immune cell composition. The quick-aging population had much more neutrophils (p-value = 9.8e-06, Wilcoxon test), less CD8 T cells (p-value = 0.041, Wilcoxon test), and less resting memory CD4 T cells (p-value = 0.017, Wilcoxon test). The decreased numbers of CD8 T cells and resting memory CD4 T cells with age were consistent with the previous studies^25,28^. Altogether, the linear-based aging clock could effectively estimate an aging rate through general model searching, capture the systematic aging change pattern to a degree, and be applied to distinguish quick- and slow-aging populations for future use.

### 5. Aging Transcriptome Undergoes a Fluctuation with a Crest around 40

From the general trajectory above (Fig. 1-B), the non-linear dimension of aging was shown yet often went unnoticed in researches with a two-group design. A recent study in human plasma proteome revealed waves of changes in the fourth, seventh, and eighth decades^10^. We wondered if there was a similar pattern at the transcriptional level. Genes with significant changes in a window period of 20 were identified by Differential Expression-Sliding Window Analysis (DE-SWAN)^10^. The algorithm takes gene expression within a window of 20 years. It compares two groups in parcels of 10 years (*e.g*., 30-40 years old compared to 40-50 years old) by routine differential expression analysis while sliding from young to old at a step size of 1 year. Gene expression changes at middle age were captured by the sliding window successively. The age distribution (Supplement Fig. 1-A) showed that the center age was restricted in the 30-60 range when analyzed from 20 to 60. The significantly changed genes around the center age with different p-value cutoffs were summarized (Fig. 4-A). Intriguingly, there was a crest at the age of 40, corresponding to the finding at the protein level. The crest remained robust at the different window sizes (Fig. S4-B, E). The genes with significant changes (p <0.05) at the age of 40 were named Wave ARGs (Supplement table 7), whose definition is different from Linear ARGs. Notably, 22 upregulated Linear ARGs were downregulated at the age of 40, and 23 genes *vice versa*. Apart from the biological processes Linear-ARGs involving in, enrichment analysis showed that (Supplement table 8) the Wave ARGs down-associated with age took part in respiratory electron transport. At the same time, the Wave ARGs up-associated with age were enriched in actin filament-based process and Rho-GTPase signaling (Fig. 4-C). Among the top Wave ARGs, *MXD1*, encoding a member of the MYC/MAX/MAD network of leucine zipper transcription factors^29^, was involved in the regulation of telomerase^30^. However, *MXD1* was not identified by linear analysis though it showed a significant upward trend around the age of 40 (adjusted p-value = 0.01, ANOVA test with sex as covariance, Benjamini-Hochberg method). Therefore, the aging transcriptome showed fluctuation with a crest around 40.

**Figure 4.**
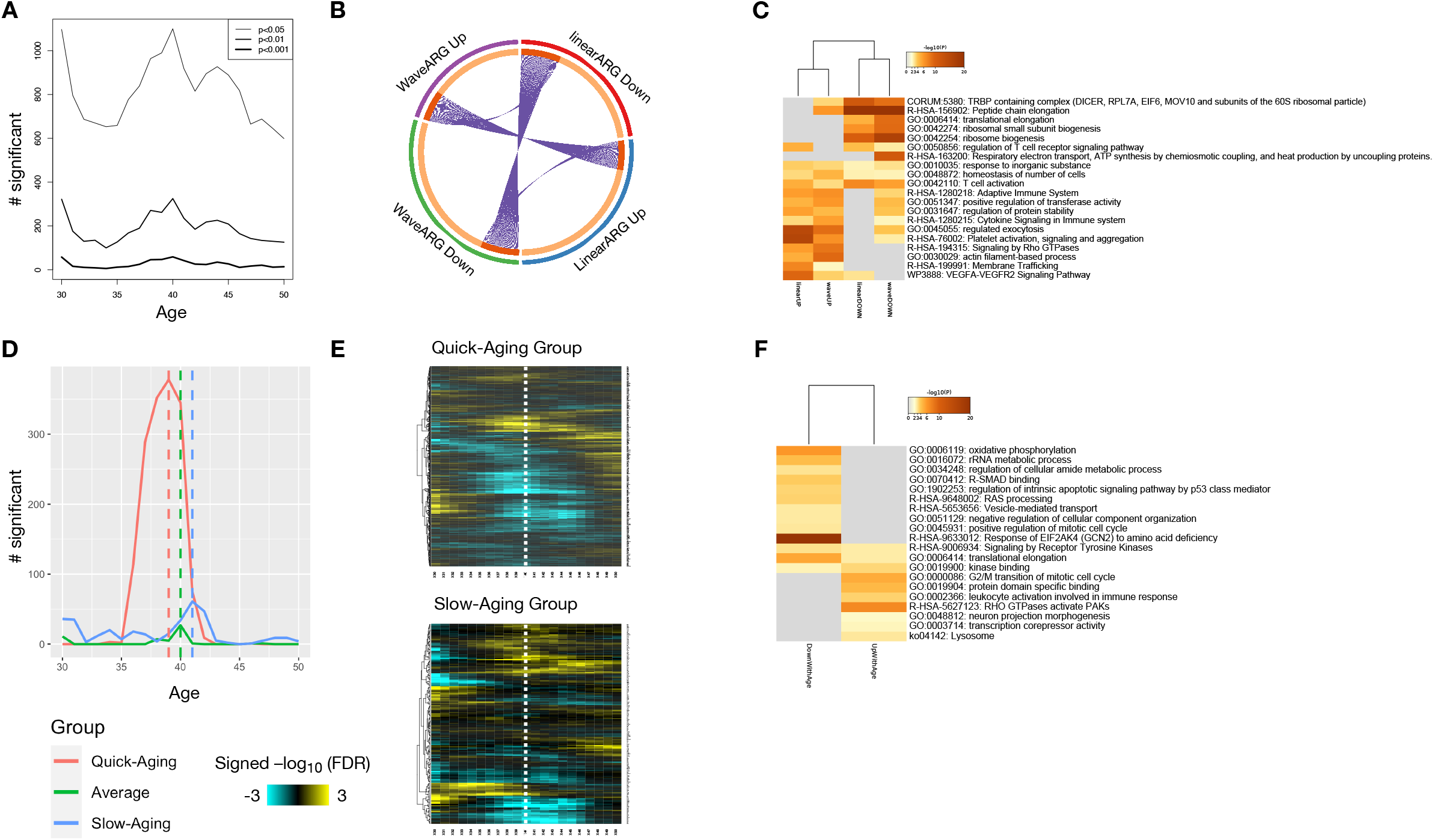
Undulating aging transcriptome with a peak around 40. (A) Count of genes with significant changes around a certain age. (p-value, ANOVA test) (B) Circus plot showing the overlap in WaveARG and LinearARG. (in both up and down direction, p-value < 0.05) (C) Heat map showing GO term enriched for WaveARG discovered in comparison with LinearARG. (D) Count of genes with significant changes peaks at different ages in groups identified by the model (i.e., Quick-Aging group, Slow-Aging group, and Average). Quick-Aging group peaks at 39, while Average peaks at 40 and Slow-Aging group peaks at 41. (q: adjusted p-value, Benjamini-Hochberg method, q<0.1) (E) Heat map showing general changes in transcriptome during aging in Quick-Aging group and Slow-Aging group. Signed −log10(FDR) were used as heat map value. The white dash line marked age 40. (F) Heat map showing GO term enriched for WaveARG in Quick-Aging Group.

### 6. Aging Transcriptome Fluctuation Differed in the Quick- and Slow-aging Populations

We then asked if the crest were different between a quick-aging population and a slow one. The same sliding window analysis was conducted on the quick-aging and slow-aging populations, respectively. In the quick-aging group, more dysregulated genes were identified at age 39, indicating that the crest came earlier and was more vigorous. However, less changed genes were found in the slow-aging group with the crest at age 41 (Fig. 4-D). The pattern remained robust with the different window sizes (Fig S4-A, C, D, F).

Furthermore, the changed magnitude between the quick-aging and the slow-aging group differed, such as the changed center shifting to the younger in the quick-aging group (Fig. 4-E). The crest in the quick-aging group was dramatic, so we conducted the enrichment analysis on these genes. The downregulated genes were involved in oxidative phosphorylation, regulation of intrinsic apoptotic pathway by p53, and the response of EIF2AK4 to amino acid deficiency. On the other hand, the upregulated genes were enriched in leukocyte activation, and RHO GTPases activate PAKs and G2/M transition of the mitotic cell cycle (Fig. 4-F, Supplement table 9). Altogether, these results showed that the aging transcriptome fluctuation at age 40 differed between the quick-aging and the slow-aging populations.

### 7. Model-based Assessment for Geroprotective Molecules

Numerous researches have been conducted in a quest for anti-aging intervention^31^. Epigenomic clocks, not the transcriptome-based ones, have been applied in the quantitative assessment of the rejuvenation effect^32^. Therefore, we applied the aging clock to assess the individual responses to star geroprotective molecules. Blood samples were collected from 8 volunteers, the same as the Method the extensive cohort above (Supplementary Fig. 5-A). Four of the collected blood samples were treated with LPS and then examined the mRNA expression of TNF compared with controls by the qPCR analysis (Supplementary Fig. 5-B). As expected, the TNF expression is significantly induced after the LPS treatment (T-paired test, p=0.005), showing that the blood samples were still responsive to external stimulations. Then, five geroprotective molecules were chosen to treat blood samples for 24 hours, including Metformin^33^, NMN^26^, Resveratrol^34^, Aspirin^35^, and Curcumin^36^, followed by the sequencing and evaluation by the aging clock after QC and pre-processing (See Method) to compare the predicted ages between treated ones and paired controls (Fig 5-A). The Metformin-treated and NMN-treated ones were predicted to be significantly younger (Metformin: 3 paired sample, t paired test: p=0.031; NMN: 6 paired sample, paired t-test: p=0.033, Fig 5-B).

**Figure 5.**
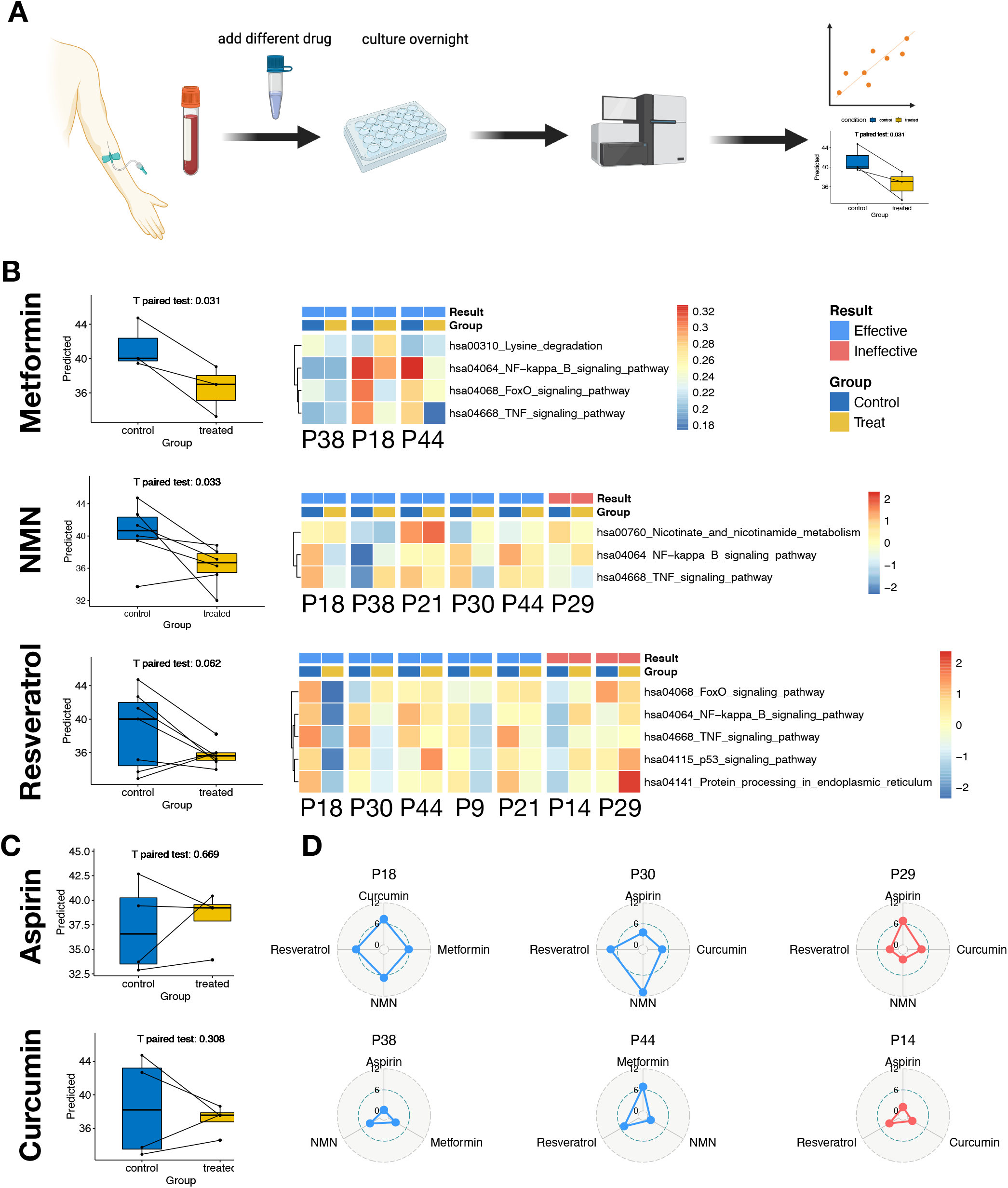
Model-based assessment for individual responses to known geroprotective molecules. (A) Graphic summary for the drug intervention and assessment pipeline. (B) Box plots show the model’s prediction of the samples treated with different molecules compared with control. (Paired t-test and the alternative hypothesis was the predicted age of control samples were less than the treated samples). Heatmaps of enrichment score of known KEGG pathways involved in specific drugs. The columns were grouped in treated and control samples of the same person (Id started with P) to see the drug effect. The drifts were further compared between individuals on whom the drug was evaluated to have an anti-aging effect or not. (C) Box plots show the model’s prediction of the samples treated with different molecules compared with control. (Paired t-test and the alternative hypothesis was the predicted age of control samples were less than the treated samples). (D) Radar charts show individual response heterogeneity to different geroprotective molecules. Blue: treated sample was predicted younger than the control. Red: treated sample was predicted older than the control. The value of radius was the predicted age difference between the treated samples and the controls.

In contrast, no significant reductions in the treated groups, using Resveratrol, Aspirin, or Curcumin, were observed (Fig. 5-B, C). The individual responses to different geroprotective molecules also differed in the predicted age-reduction scale. For example, the blood sample (#P44) responded best to Metformin while the #P30 one showed the best response to NMN while P14 and P29 ones responded poorly to all treated molecules (Fig. 5-D). KEGG^37^ pathway enrichment scores were calculated for each sample, and the drifts between control and treated samples were observed. The drifts were further compared among individuals to evaluate the anti-aging effect. The Metformin-treated samples generally had a higher enrichment in Lysine degradation (KDAC) pathway and lower enrichments in nuclear factor-kB (NF-kB), forkhead box transcription factors (FOXO), and tumor necrosis factor (TNF) pathways, in agreement with its molecular mechanism in aging^38^. The Nicotinate and nicotinamide metabolism pathways were generally augmented in NMN-treated samples, accompanied by decreased enrichment of NF-kB and TNF pathways. The Resveratrol-treated samples with the effective responses had an attenuated enrichment in FOXO, NF-kB, TNF, p53, and protein-processing in endoplasmic reticulum pathways^39^, while the ineffective ones showed an opposite drift (Fig. 5-B). The enriched 5-monophosphate-activated protein kinase (AMPK) and oxidative phosphorylation pathways were identified in Aspirin-treated samples of the affected group. The curcumin-treated samples showed decreased enrichment in mammalian target of rapamycin (mTOR), FOXO, transforming growth factor β (TGF-β), NF-kB, and TNF pathways, while samples in the ineffective group did not (Supplementary Fig 5-C). These results provided a molecular view for the different individual responses, and the aging clock predicted a younger age after the Metformin- and NMN-treated samples.

## Discussion

The study deeply mined the aging blood transcriptome, revealed the underlying midlife change in gene expression, and constructed an ensemble aging clock and a more straightforward linear aging clock, which shows the promising application in predicting personalized drug rejuvenation effect.

This study identified a pool of age-related genes. It should be noted that these Linear ARGs presented in this study depend on this cohort, which is the genetic background of the Eastern Chinese population. A meta-analysis demonstrated that the aging transcriptome signature displayed low overlap in different genetic backgrounds, such as native, Hispanic, and African American^6^. Although a universal aging pattern is desirable, aging biomarkers specific to a particular genetic background population should be studied. Therefore, the Linear ARGs, together with gender, were used as features for the aging clock. A study showed that the aging biomarkers were population-specific for South Korean, Canadian, and Eastern European so that aging clocks for each population were built up^40^. Although the ethnic background should be considered when constructing aging clocks, part of the aging pattern and the general methodology should be consistent.

An ensemble LDA model was built in a recent study^8^ and showed better performance in an age bin approach, indicating the ensemble model is a promising structure. For a complex process with linear and non-linear changes, such as aging, a general ensemble model combing linear and non-linear models is a suitable structure. In this study, the ensemble model showed a slight advantage in accuracy compared to the elastic net-based model. However, for simplicity in interpretation and application, the elastic net-based linear model was chosen. This may be due to the relatively small cohort size (505) and sample distribution. Although the cohort covered an age range of 18 to 68 and the median age of 36, the old samples were insufficient. Therefore, a more extensive and comprehensive cohort is necessary for future study.

The division of quick- and slow-aging populations was clinically meaningful for risk evaluation, treatment, and personalized anti-aging therapy. The immune cell composition of the quick-aging group shift toward an older phenotype. Neutrophils-Lymphocyte ratio (NLR) is a well-accepted marker for systematic inflammation related to the prognosis of cancer^41^, cardiovascular diseases, and all-cause mortality^42^. The quick-aging group displayed higher Neutrophils and lower Lymphocyte count, indicating a higher degree of systematic inflammation. Transcriptome-based aging clocks have an advantage in interpretability. Thus, it can be used for long-term monitoring, such as physical examination, and provide other information along with the aging clock.

In the non-linear dimension of aging, the patterns of gene expression undergo dynamic changes throughout life. The fluctuation should be considered when gene signatures are for diagnostic purposes, improving the specificity and accuracy. Modules mapping gene changes (Fig. 1) were associated with the hallmark activities in aging. The gene expression variances in life suggested the role of environmental factors, mental health^43^, and other soft factors apart from the genetic programming (the hard factor) in aging, especially in the midlife change. The quick-aging group showed an earlier and more prominent for the aging change. However, our findings showed that early anti-aging interventions in midlife, more investigations on these age-related Wave genes of the quick-aging group aid in dissecting the heterogeneous aging process.

The aging clock succeeded in evaluating the rejuvenation effect of molecules such as NMN and Metformin *in vitro*. The aging clock was applied to control and paired drug-treated samples to get the relative age prediction. The method can be applied for *in vitro* screening for anti-aging interventions. Most treated blood samples of responsive individuals showed the enriched signaling pathways involved in the molecular mechanism related to aging. Consistently, the treated ones with the poorly responses showed the opposite enrichment, displaying the enriched inflammatory pathways.

Moreover, these results indicated that each person could respond differently to each molecule, as their responses to the molecular targets and the related mechanisms vary. Thus, the aging clock can be used to determining the most suitable drugs. However, these observations may need to be further validated considering the limited samples. Therefore, we aim to develop an aging clock by investigating the transcriptome landscape of age-related diseases on more samples in the future.

## Methods

### Blood Sample Acquisition and RNA-seq

Blood samples were drawn from people coming for physical examination. Approval for utilizing the samples was obtained from the Ethics Committee of the Second Affiliated Hospital, School of Medicine, Zhejiang University (Approval Reference Number: 2019-234). Next, samples were first treated with ACK Lysis Buffer (Solarbio, China). Samples were processed for RNA-seq, which was modified from a previous method^44^. Blood samples were first lysed by Trizol reagent (TAKARA). Then, reverse transcription was conducted using SuperScript II reverse transcriptase (Invitrogen), and double-strand cDNA was synthesized using NEBNext mRNA second strand synthesis kit (NEB). Cleaning was done using AMPure XP beads (Beckman Coulter), and the sequencing library was constructed using the Nextera XT kit(Illumina). The pooled library was sequenced on the Illumina X-Ten platform. RNA-seq reads data were mapped to the reference genome using STAR^45^. Expression was calculated with counts per million (CPM).

### Cell Culture

Whole blood samples from 8 individuals were randomly selected for treatment and culture. Each sample was divided into six portions (100 μl each, some samples had less than six due to the limit volume of blood) and were added to the 48-well plate (Supplementary Fig 5-A). Six replicates of each person were added different reagents at reported concentrations (100*μ*M Aspirin^46^(Selleck, S3017), 50*μ*g/ml Curcumin^47^(Selleck, S1039), 50*μ*M Resveratrol^48^(Selleck, S1396), 500*μ*M NMN^49^ (Qingyuan Shengyi Biological Technology Co., Ltd.), 100*μ*M Metformin^50^(Selleck, S1950), 100ng/ml LPS(Sigma, L2880) respectively. Then these samples were incubated and constantly rotated on a shaker at 6 rpm, 37°C for 24 hours^51^. Then these samples were harvested and washed with ACK lysis buffer (Solarbio, China) three times to remove the erythrocytes before RNA-seq mentioned above.

### Data Quality Control

Samples with total CPM three times the mean absolute deviation higher or lower away from the medium were filtered. For the large cohort, samples with age three times the mean absolute deviation higher or lower away from the medium were filtered. Moreover, we only kept genes that were expressed in at least 10% of all samples. Supplementary Fig 1-B, C showed the library size and number of the expressed gene of the large cohort.

### Enrichment Analysis

To determine the biological meaning of a group of genes, we queried GO and Reactome terms using Metascape^18^.

### Clustering of Gene Expression Trajectories

To estimate trajectories of age-related genes (roughly selected by person correlation score > 0.05) during aging, the expression trajectories of 4318 genes are fitted by loess. To reduce the complexity in changing patterns, the trajectories were clustered by unsupervised hierarchical clustering. Genes with similar changing patterns were poured into the same module. To understand the biological functions of each cluster, we queried GO databases, using R clusterProfiler package^52^ and org.Hs.eg.db package^53^.

### Linear Fitting and Linear ARGs of the Blood Transcriptome

Linear fitting was done by glm function in the R stats package and gaussian family. For each gene, the linear model fits as follow:

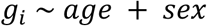

The square sum of was calculated by the aov function in the R stats package. The percentages of variance explained by sex and age for each gene were computed in the form of:

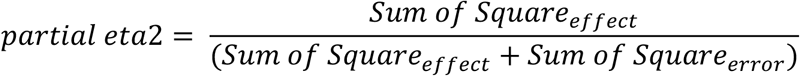

The age effect for each gene was determined by the two-side p-value of t-test in summary.lm function in R stats package. Genes with a significant age effect (p-value >0.05) were considered as Linear ARGs.

### Construction of Aging Clock

AutoGluon^20^ is an open-source auto-machine learning framework, and the AutoGluon-Tabular, which was designed for structured data, was applied for model searching (python 3.8.5, autogluon 0.2.0). The cohort was first divided as train set and test set with a ratio of 3:1, and then the train set was further divided, 80% of which was used for model construction and 20% of which (validation set) was used for model validation in a search for the best model. The train and test set were separately scaled and centered in preprocessing step. 1138 Linear ARGs and gender were used as model features. The models were trained in MAE (mean absolute error) and tested in MAE and other metrics. The hyperparameters space was expanded from default and stated in the supplementary data.

The Elastic-net model was built in R (4.0.5) by the glmnet package. The cv.glmnet function was used for the parameter lambda search with 20 fold cross-validation and MAE as measuring metric. An outer loop of 10-fold cross-validation was applied for an average MAE. Parameter alpha was determined by grid search and the best average MAE. The final model was constructed by the best alpha with correspondent lambda.

### Aging rate and Quick/Slow Aging Population Distinguishment

Aging clock prediction was used for aging rate estimation and was calculated with the difference between the model predicted age and chronological age:

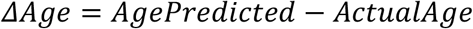

Considering the prediction error distribution, it was adjusted for age itself, and the data set difference. (0-train set, 1-test set)

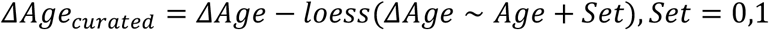

With the curated aging rate, the population was further classified into average, quick-aging, and slow-aging groups. *q*_1_, *q*_2_, *q*_3_ were the ascending quantile numbers of the curated aging rate of the cohort. People with Δ*Age_curated_* > *q*_3_ was classified into the quick-aging group and people with Δ*Age_curated_* < *q*_1_ were classified into slow-aging group.

### Sliding Window Analysis

DE-SWAN^10^ was used with gender as covariant, and the bin size of 10 and 15 was tested. The number of significantly changed gene in the window were summarized.

### Model Assessment in Cultured Sample

The gene expression matrix was first scaled and centered to gaussian distribution. Then the gender information was appended. The samples were predicted by the elastic-net-based model. The predicted ages of the treated and control sample were compared by paired t-test.

### KEGG Pathway Enrichment Analysis

The gsva function of R package GSVA^56^ was utilized with parameters as follows: min.sz of 5, max.sz of 500, “ssgsea” method, abs.ranking and other default parameters. The KEGG gene sets were obtained from the KEGG pathway database of release 99.1.

**Figure 6.**
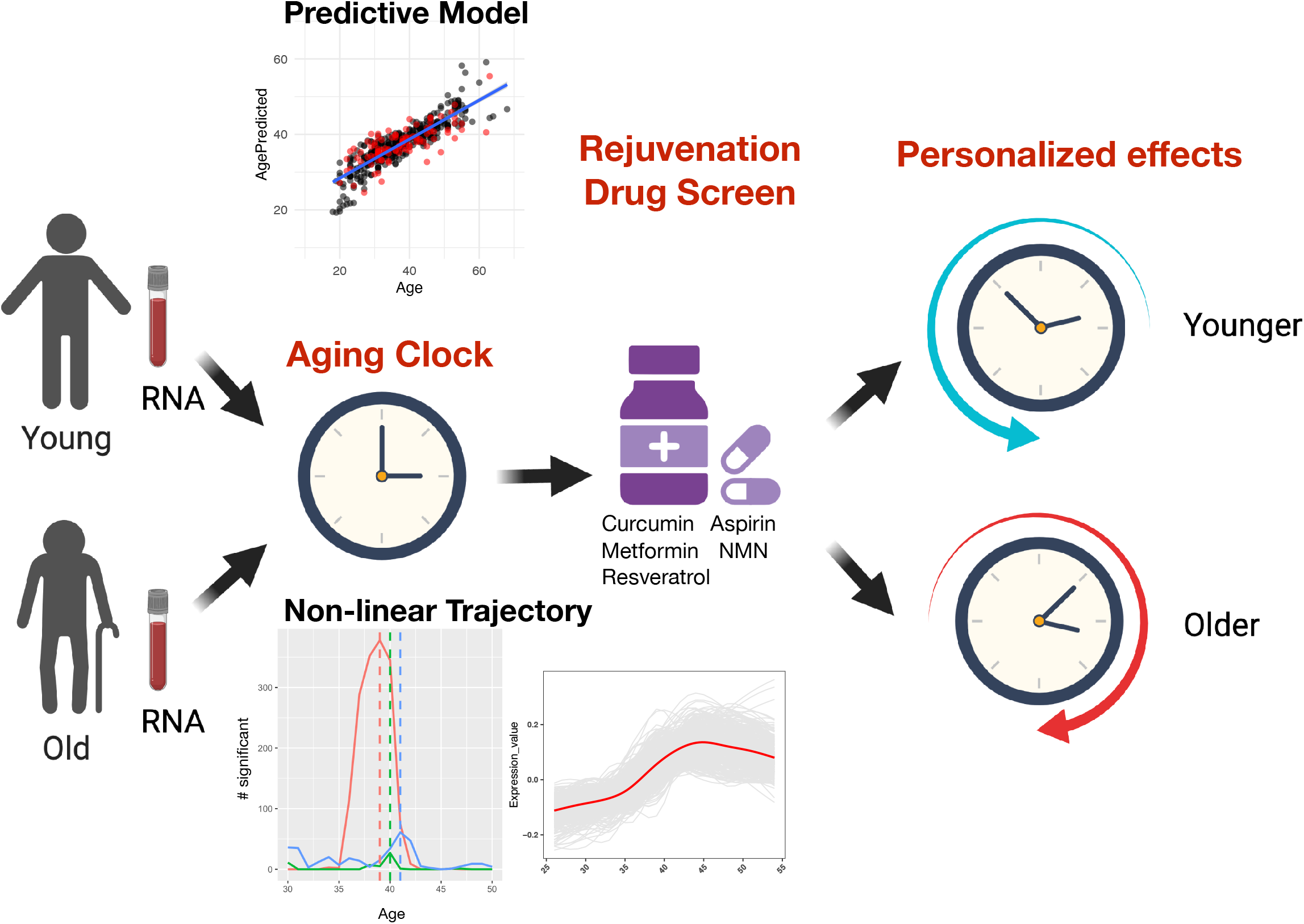
Graphical Summary.

## Data Availability

All the data generated in this study are available upon reasonable request to the corresponding author.

## Code Availability

All the codes generated in this study are available upon reasonable request to the corresponding author.

## Author Contributions

X.S.: Study design, data analysis, sample processing and manuscript writing; B.W.: Study design, acquisition of clinical sample, sample processing; W.J.: Data analysis, sample processing and manuscript writing; Y.L., N.N., L.G., Y.L.: Sample processing; Y.Z.: Acquisition of clinical sample; K.Z.: Data analysis; J.L.: Manuscript revision; J.J.: Acquisition of clinical sample and design; X.Z., H.O.: Conception and design.

## Acknowledgement

This work was supported by National Key R&D Program of China (2017YFA0104900);National Natural Science Foundation of China (81871127, 8217060567, 31870973); Zhejiang Medical and Health Science and Technology Program (2013KYB080); Science and Technology program of Jinhua Science and Technology Bureau (Grant No. 2021-3-001)

## Supplementary data legend

**Figure S1.**
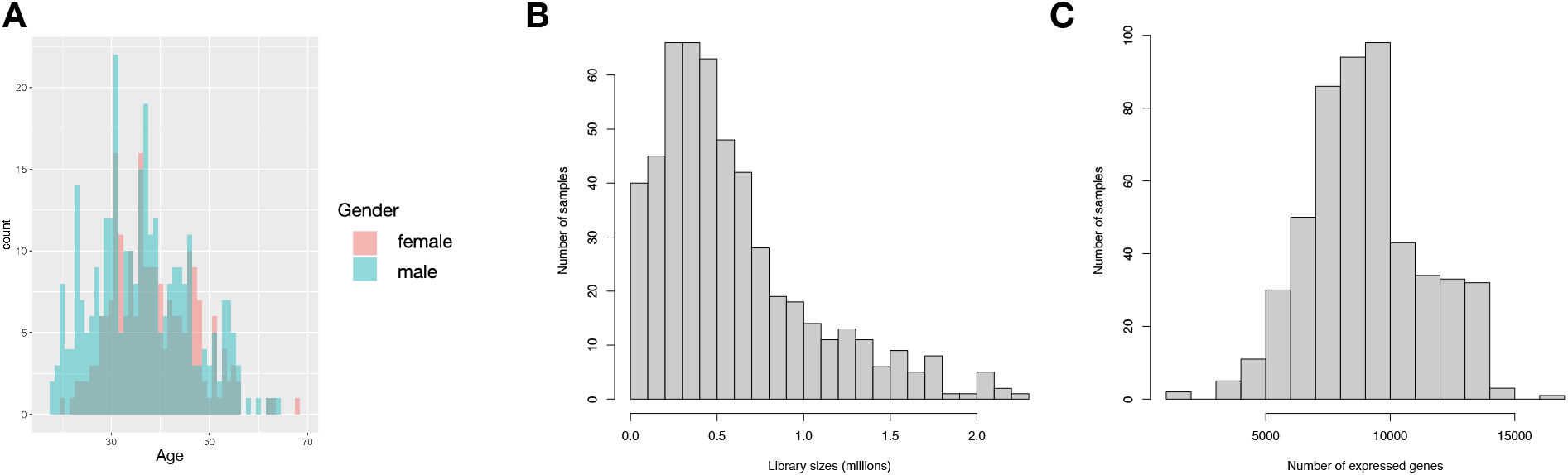
Cohort characterization of Chinese population across the life span. (A) Histogram shows sample distribution in this cohort. (B) Histogram shows sample’s library size distribution of the RNA sequencing data. (C) Histogram shows sample’s number of expressed genes distribution of the RNA sequencing data.

**Figure S2.**
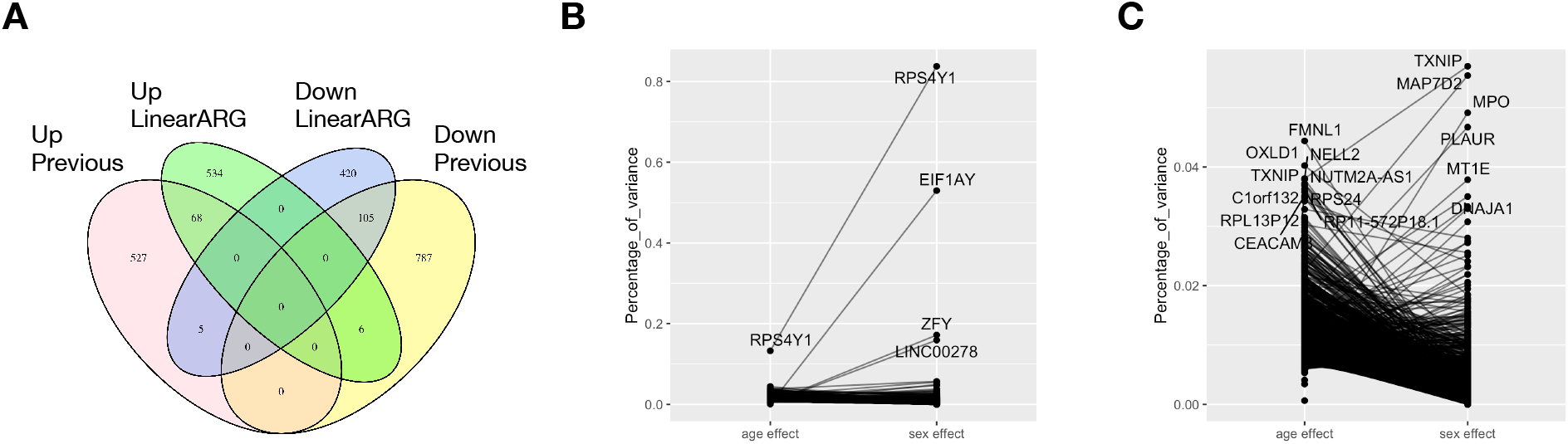
The association of Linear Age-related Genes (Linear ARGs) with previous studies and sex. (A) Venn plot of LinearARGs and genes reported in the previous meta-study with direction. (B-C) Percentage of variance explained by age and sex in the linear fitting. (ANOVA test, Sum of square)

**Figure S3.**
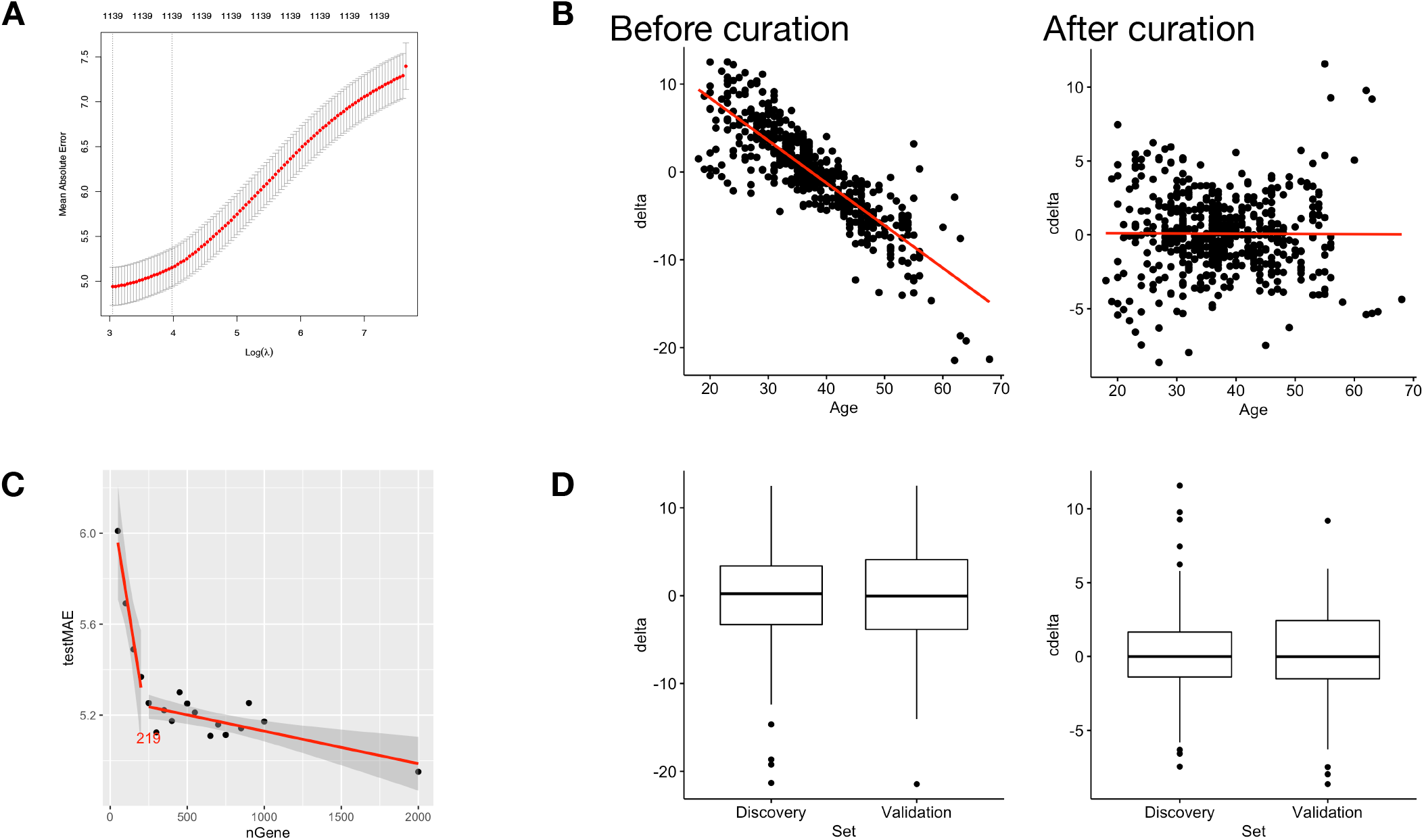
Characterization of Aging Clock modeling. (A) Relationship between lambda and train error for elastic-net modeling. In this study, lambda of minimum mean absolute error (MAE) is picked. (B) The correlation between actual age and delta-age in a model before and after model curation for age. (C) Relationship between MAE in the test set and the number of genes in the model, of which 219 is the turning point. (Genes are selected according to the rank of p-value in figure 2) (D) The difference of delta-age between training and testing data set in a model before (left) and after (right) model curation for data set.

**Figure S4.**
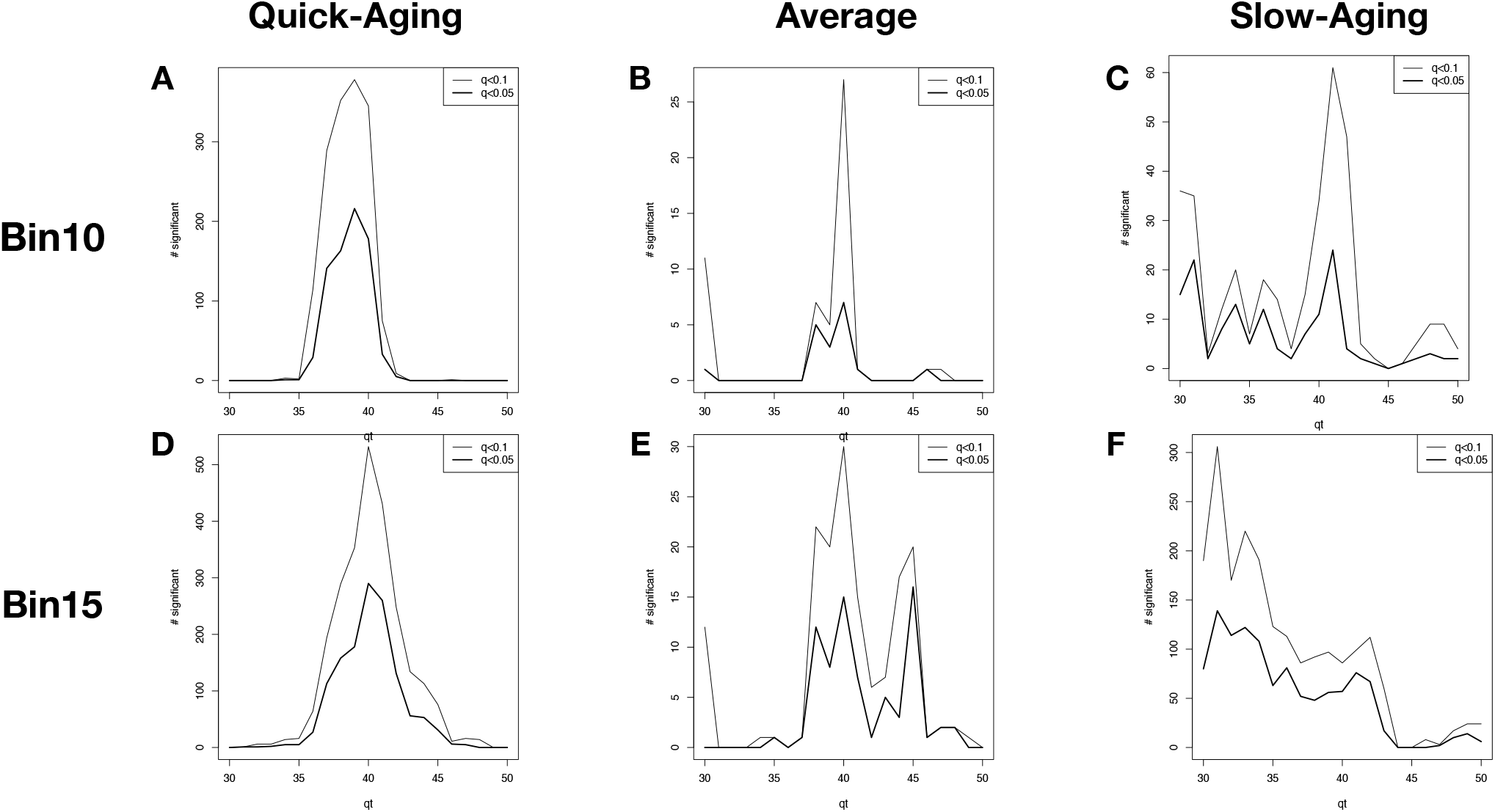
Wave ARGs distribution in different window sizes across the group. (A-C) Changes in the number of genes experienced significant changes in a window size of 20 (bin size is half of the window size, thus window size of 20, the bin size is 10. The figures show gene changes in −10 ~ +10 around center age). (D-F) Changes in the number of genes experienced significant changes in a window size of 30 (−15 ~ +15).

**Figure S5.**
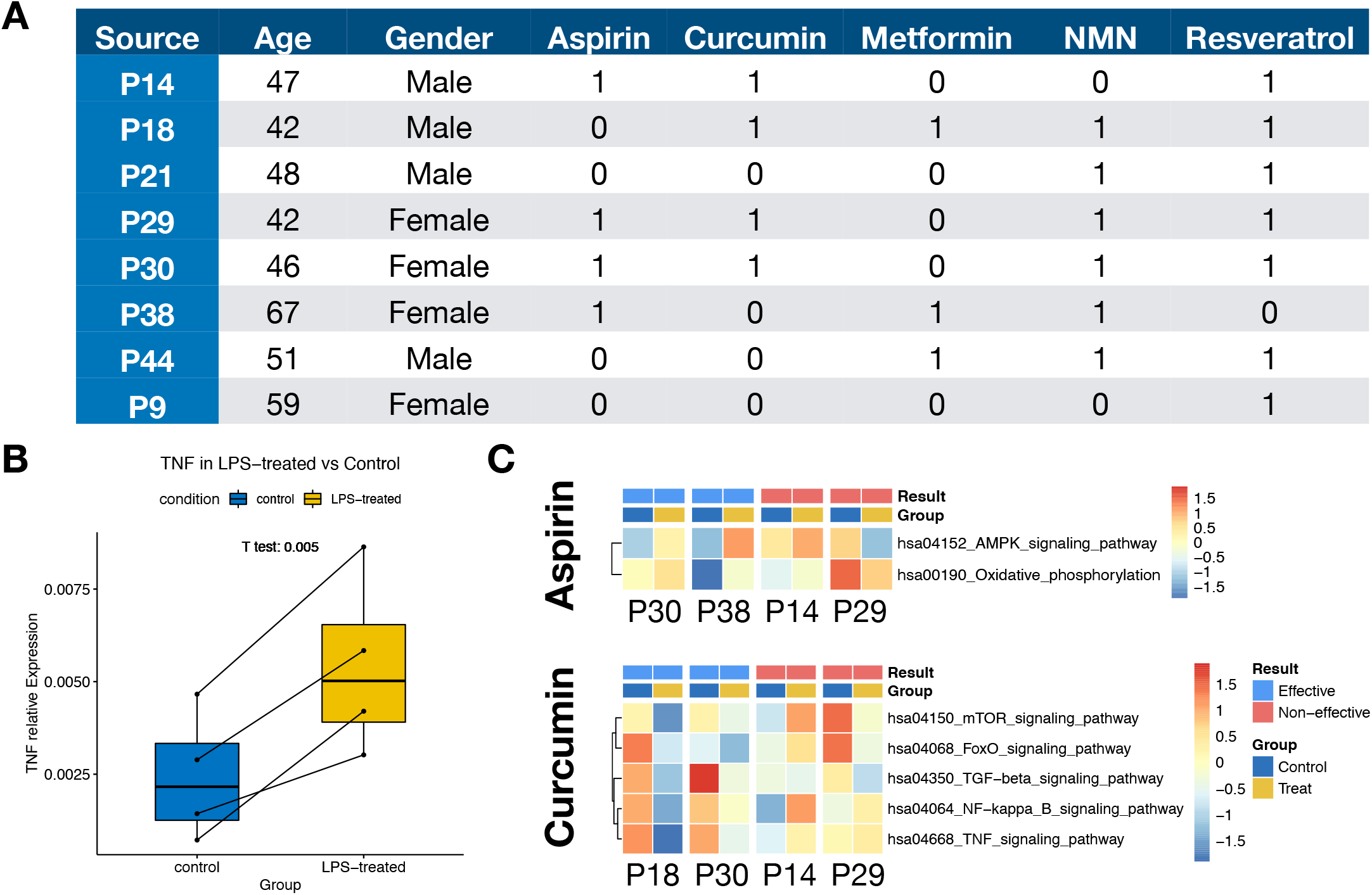
Characterization of molecule-treated samples. (A) Table of the existed sample. (1 - sample existed, 0 - sample not existed). The sample not existed is lost due to technical error. (B) Box plot shows the qPCR relative expression of TNF in LPS treated sample in comparison with control. (Paired t-test) (C) Heatmap of enrichment score of known KEGG pathways involved in Aspirin and Curcumin. The columns were grouped in treated and control samples of the same person (Id started with P) to see the drug effect. The drifts were further compared between individuals on whom the drug was evaluated to have an anti-aging effect or not.

## Supplement table legend

**Supplement table 1. Gene ontologies of eight aging trajectories;**

**Supplement table 2. List of linear ARGs;**

**Supplement table 3. Gene ontologies of linear ARGs;**

**Supplement table 4. Model parameter and performance metrics of autogluon;**

**Supplement table 5. Feature importance of the ensemble model inferred by autogluon;**

**Supplement table 6. Parameter grid search result for the elastic-net model;**

**Supplement table 7. List of Wave ARGs;**

**Supplement table 8. Gene ontologies of linear and wave ARGs in comparison;**

**Supplement table 9. Gene ontologies of wave ARGs in quick aging group.**

## References

1. United Nations, Department of Economic and Social Affairs, Population Division (2019). World Population Prospects 2019: Highlights. ST/ESA/SER.A/423.

2. López-Otín, C., Blasco, M. A., Partridge, L., Serrano, M. & Kroemer, G. The hallmarks of aging. Cell 153, 1194–1217 (2013).

3. Alpert, A. et al. A clinically meaningful metric of immune age derived from high-dimensional longitudinal monitoring. Nat. Med. 25, 487–495 (2019).

4. Sayed, N. et al. An inflammatory aging clock (iAge) based on deep learning tracks multimorbidity, immunosenescence, frailty and cardiovascular aging. Nat. Aging 1, 598–615 (2021).

5. Valdes, A. M., Glass, D. & Spector, T. D. Omics technologies and the study of human ageing. Nat. Rev. Genet. 14, 601–607 (2013).

6. NABEC/UKBEC Consortium et al. The transcriptional landscape of age in human peripheral blood. Nat. Commun. 6, 8570 (2015).

7. Mamoshina, P. et al. Machine Learning on Human Muscle Transcriptomic Data for Biomarker Discovery and Tissue-Specific Drug Target Identification. Front. Genet. 9, 242 (2018).

8. Fleischer, J. G. et al. Predicting age from the transcriptome of human dermal fibroblasts. Genome Biol. 19, 221 (2018).

9. Xia, X., Wang, Y., Yu, Z., Chen, J. & Han, J.-D. J. Assessing the rate of aging to monitor aging itself. Ageing Res. Rev. 69, 101350 (2021).

10. Lehallier, B. et al. Undulating changes in human plasma proteome profiles across the lifespan. Nat. Med. 25, 1843–1850 (2019).

11. Elliott, M. L. et al. Disparities in the pace of biological aging among midlife adults of the same chronological age have implications for future frailty risk and policy. Nat. Aging 1, 295–308 (2021).

12. Kevei, É. & Hoppe, T. Ubiquitin sets the timer: impacts on aging and longevity. Nat. Struct. Mol. Biol. 21, 290–292 (2014).

13. Márquez, E. J. et al. Sexual-dimorphism in human immune system aging. Nat. Commun. 11, 751 (2020).

14. Kauppila, T. E. S., Kauppila, J. H. K. & Larsson, N.-G. Mammalian Mitochondria and Aging: An Update. Cell Metab. 25, 57–71 (2017).

15. Turi, Z., Lacey, M., Mistrik, M. & Moudry, P. Impaired ribosome biogenesis: mechanisms and relevance to cancer and aging. Aging 11, 2512–2540 (2019).

16. Austad, S. N. & Fischer, K. E. Sex Differences in Lifespan. Cell Metab. 23, 1022–1033 (2016).

17. Zhang, W. et al. A single-cell transcriptomic landscape of primate arterial aging. Nat. Commun. 11, 2202 (2020).

18. Zhou, Y. et al. Metascape provides a biologist-oriented resource for the analysis of systems-level datasets. Nat. Commun. 10, 1523 (2019).

19. Golomb, L., Volarevic, S. & Oren, M. p53 and ribosome biogenesis stress: The essentials. FEBS Lett. 588, 2571–2579 (2014).

20. Erickson, N. et al. AutoGluon-Tabular: Robust and Accurate AutoML for Structured Data. ArXiv200306505 Cs Stat (2020).

21. Caruana, R., Niculescu-Mizil, A., Crew, G. & Ksikes, A. Ensemble selection from libraries of models. in Twenty-first international conference on Machine learning - ICML ‘04 18 (ACM Press, 2004). doi:10.1145/1015330.1015432.

22. Hilaire, C. S. et al. NT5E Mutations and Arterial Calcifications. N Engl J Med 11 (2011).

23. Su, S. et al. Proteomic Analysis of Human Age-related Nuclear Cataracts and Normal Lens Nuclei. Investig. Opthalmology Vis. Sci. 52, 4182 (2011).

24. Xia, X. et al. Three-dimensional facial-image analysis to predict heterogeneity of the human ageing rate and the impact of lifestyle. Nat. Metab. 2, 946–957 (2020).

25. Qin, L. et al. Aging of immune system: immune signature from peripheral blood lymphocyte subsets in 1068 healthy adults. Aging 8, 848–859 (2016).

26. Newman, A. M. et al. Determining cell type abundance and expression from bulk tissues with digital cytometry. Nat. Biotechnol. 37, 773–782 (2019).

27. Newman, A. M. et al. Robust enumeration of cell subsets from tissue expression profiles. Nat. Methods 12, 453–457 (2015).

28. Zheng, Y. A human circulating immune cell landscape in aging and COVID-19. 31 (2020).

29. Cascón, A. & Robledo, M. MAX and MYC: A Heritable Breakup: Figure 1. Cancer Res. 72, 3119–3124 (2012).

30. Schaefer, C. F. et al. PID: the Pathway Interaction Database. Nucleic Acids Res. 37, D674–D679 (2009).

31. Partridge, L., Fuentealba, M. & Kennedy, B. K. The quest to slow ageing through drug discovery. Nat. Rev. Drug Discov. 19, 513–532 (2020).

32. Fahy, G. M. et al. Reversal of epigenetic aging and immunosenescent trends in humans. Aging Cell 18, (2019).

33. Cabreiro, F. et al. Metformin Retards Aging in C. elegans by Altering Microbial Folate and Methionine Metabolism. 12.

34. Baur, J. A. & Sinclair, D. A. Therapeutic potential of resveratrol: the in vivo evidence. DRUG Discov. 14.

35. McNeil, J. J. et al. Effect of Aspirin on Cardiovascular Events and Bleeding in the Healthy Elderly. N. Engl. J. Med. 379, 1509–1518 (2018).

36. Zia, A., Farkhondeh, T., Pourbagher-Shahri, A. M. & Samarghandian, S. The role of curcumin in aging and senescence: Molecular mechanisms. Biomed. Pharmacother. 134, 111119 (2021).

37. Kanehisa, M., Furumichi, M., Sato, Y., Ishiguro-Watanabe, M. & Tanabe, M. KEGG: integrating viruses and cellular organisms. Nucleic Acids Res. 49, D545–D551 (2021).

38. Torres, W. et al. Anti-Aging Effect of Metformin: A Molecular and Therapeutical Perspective. Curr. Pharm. Des. 26, 4496–4508 (2020).

39. Zhou, D.-D. et al. Effects and Mechanisms of Resveratrol on Aging and Age-Related Diseases. Oxid. Med. Cell. Longev. 2021, 1–15 (2021).

40. Mamoshina, P. et al. Population Specific Biomarkers of Human Aging: A Big Data Study Using South Korean, Canadian, and Eastern European Patient Populations. J. Gerontol. Ser. A 73, 1482–1490 (2018).

41. Templeton, A. J. et al. Prognostic Role of Neutrophil-to-Lymphocyte Ratio in Solid Tumors: A Systematic Review and Meta-Analysis. JNCI J. Natl. Cancer Inst. 106, (2014).

42. Wang, X. et al. Neutrophil to lymphocyte ratio in relation to risk of all-cause mortality and cardiovascular events among patients undergoing angiography or cardiac revascularization: A meta-analysis of observational studies. Atherosclerosis 234, 206–213 (2014).

43. Wertz, J. et al. Association of History of Psychopathology With Accelerated Aging at Midlife. JAMA Psychiatry 78, 530 (2021).

44. Picelli, S. et al. Smart-seq2 for sensitive full-length transcriptome profiling in single cells. Nat. Methods 10, 1096–1098 (2013).

45. Dobin, A. et al. STAR: ultrafast universal RNA-seq aligner. Bioinforma. Oxf. Engl. 29, 15–21 (2013).

46. Vitale, P. et al. Synthesis, Pharmacological Characterization, and Docking Analysis of a Novel Family of Diarylisoxazoles as Highly Selective Cyclooxygenase-1 (COX-1) Inhibitors. J. Med. Chem. 56, 4277–4299 (2013).

47. Li, N. et al. Curcumin and Curcumol Inhibit NF-*κ* B and TGF-*β*_1_/Smads Signaling Pathways in CSE-Treated RAW246.7 Cells. Evid. Based Complement. Alternat. Med. 2019, 1–9 (2019).

48. Mai, A. et al. Study of 1,4-Dihydropyridine Structural Scaffold: Discovery of Novel Sirtuin Activators and Inhibitors. J. Med. Chem. 52, 5496–5504 (2009).

49. Penke, M. et al. Oleate ameliorates palmitate-induced reduction of NAMPT activity and NAD levels in primary human hepatocytes and hepatocarcinoma cells. Lipids Health Dis. 16, 191 (2017).

50. Suh, D. H., Lee, S., Park, H.-S. & Park, N. H. Medroxyprogesterone Reverses Tolerable Dose Metformin-Induced Inhibition of Invasion via Matrix Metallopeptidase-9 and Transforming Growth Factor-β1 in KLE Endometrial Cancer Cells. J. Clin. Med. 9, 3585 (2020).

51. Orecchioni, M. et al. Single-cell mass cytometry and transcriptome profiling reveal the impact of graphene on human immune cells. Nat. Commun. 8, 1109 (2017).

52. Yu, G., Wang, L.-G., Han, Y. & He, Q.-Y. clusterProfiler: an R package for comparing biological themes among gene clusters. Omics J. Integr. Biol. 16, 284–287 (2012).

53. Carlson, M. org.Hs.eg.db: Genome wide annotation for Human. (2019) doi:10.18129/B9.bioc.org.Hs.eg.db.

